# Identification and transcriptional regulation of the mir-218-1 alternative promoter

**DOI:** 10.1101/2020.09.29.319202

**Authors:** Brenna A. Rheinheimer, Lukas Vrba, Bernard W Futscher, Ronald L Heimark

## Abstract

**Background:** miRNAs are small, endogenous non-coding RNAs approximately 22 nucleotides in length that account for approximately 1% of the genome and play key regulatory roles in multiple signaling pathways. mir-218-1 is an intronic miRNA located within intron 15 of the *SLIT2* gene. Public datasets showed enrichment of H3K4me3 within intron 4 of the SLIT2 gene. Therefore, we sought to determine the genomic location and transcriptional regulatory elements of the mir-218-1 candidate alternative promoter in pancreatic ductal adenocarcinoma.

**Methods:** Expression of mir-218 was evaluated in a panel of pancreatic ductal adenocarcinoma cell lines. The mir-218-1 candidate alternative promoter was characterized by chromatin immunoprecipitation, Sequenom, and luciferase assays. Transcriptional regulation of the mir-218-1 candidate alternative promoter was assessed using chromatin immunoprecipitation and an inhibitor to NF-kB.

**Results:** We found that expression of mir-218-1 does not correlate with SLIT2 expression and that mir-218-1 has a novel transcriptional start site separate from the SLIT2 promoter. This novel transcriptional start site showed transcriptional activity and was regulated by NF-kB.

**Conclusions:** mir-218-1 is transcribed from an independent and novel transcriptional start site located within intron 4 of the SLIT2 gene in pancreatic ductal adenocarcinoma. Additionally, mir-218-1 expression is regulated by Nf-kB at this alternative transcriptional start site in pancreatic cancer.

## Introduction

miRNAs are small endogenous non-coding RNAs approximately 22 nucleotides in length that account for approximately 1% of the genome and play key regulatory roles in multiple signaling pathways^123^. Pancreatic cancer is no stranger to miRNA dysregulation during every stage of the disease—including invasion and metastasis. Li et al. reported lower expression of miR-146a in pancreatic cancer cells compared to immortalized human pancreatic ductal epithelium (HPDE) and re-expression of miR-146a in pancreatic cancer cells inhibited their invasive capability^259^. Hypermethylation-mediated silencing of miR-124 was associated with poor survival of pancreatic ductal adenocarcinoma patients while functional studies showed that miR-124 inhibited pancreatic cancer cell proliferation, invasion and metastasis^260^. Increased miR-10b expression in pancreatic ductal adenocarcinoma is a marker of disease aggressiveness that can be measured in the plasma of patients^261^. Additionally, Ouyang et al. demonstrated that overexpression of miR-10b increased pancreatic cancer cell proliferation and tumor growth in an orthotopic model. Approximately one quarter of miRNAs are found within an intron of a coding gene implying that they are not transcribed from their own promoters^125–127^; however, it has recently been shown that roughly one-third of intronic miRNAs are transcribed from their own transcriptional units separate from their host gene^262^. A model was recently proposed suggesting that the transcription of intronic microRNAs largely coincide with the expression of their host gene. In this model, Drosha cleaves the pre-miRNA before the intron is spliced out and the two host gene exons are joined together^263^.

mir-218-1 is an intronic miRNA located within intron 15 of the *SLIT2* gene. Additionally, mir-218-2 is an intronic miRNA located within intron 14 of the *SLIT3* gene. The mature sequence for mir-218-1 and mir-218-2, termed miR-218, is identical. The differences in sequence between the two isoforms of miR-218 are located at the 3’ end of the individual stem loop structures. The gene organization and protein structure between *SLIT2* and *SLIT3* are very similar as well as the location of the mir-218-1 and mir-218-2 stem loops. Although their genomic locations are different—*SLIT2* is located on chromosome 4p15.2 and *SLIT3* is located on chromosome 5q34-35—according to the Human Protein Atlas, their expression levels and patterns in normal tissue is parallel. Epigenetic regulation of *SLIT2* and *SLIT3* by DNA methylation of their respective CpG islands in breast, lung, colorectal, nasopharyngeal, and pancreatic cancers has also been reported^180,205,206,253,264^. Several publications have also insinuated that mir-218-1 and mir-218-2 are transcribed along with their host genes *SLIT2* and *SLIT3*, respectively^179,180,182^. Since my expression data indicated that mir-218-1 is present in pancreatic ductal adenocarcinoma cell lines that lack *SLIT2* transcripts, I proposed that there is an additional transcriptional start site for mir-218-1 separate from the *SLIT2* transcriptional start site. Therefore, I sought to determine the genomic location and transcriptional regulatory elements of the mir-218-1 candidate alternative promoter in pancreatic ductal adenocarcinoma.

Along with controlling transcriptional regulation, histone modifications can also identify transcriptional starts sites within the genome. Approximately 1-2% of the human genome is located within 500 base pairs of either side of a transcriptional start site, yet almost 25% of the genome is potentially transcribable^265^. It has been shown that methylation on H3K4 and acetylation on H3K9 and H3K14 are localized to the 5’ regions of transcriptionally active genes, but decreased downstream of transcriptional start sites^265^ suggesting that H3K4 methylation and H3K9/H3K14 acetylation may be required for transcription initiation and transition to elongation, but do not track with RNA polymerase II as it moves throughout the transcribed region. Instead, large transcribed regions of human genes are kept in a deacetylated conformation allowing elongating polymerase to read and transcribe these regions^227^. Interestingly, several years later, it was discovered that specific histone modifications could identify gene promoters, enhancers, silencers, and insulators. In fact, H3K4me3 is considered a hallmark of actively transcribed promoters in eukaryotes^266^. Several studies have also shown that novel promoters for protein-coding genes can be identified on the basis of H3K4me3 enrichment^267–269^.

Considering that miRNAs use the same transcriptional machinery as protein-coding genes, it was no surprise that histone modification H3K4me3 could also be used to precisely map the locations of miRNA transcriptional start sites^262^. Ozsolak et al. elucidated the transcription initiating regions of 175 miRNAs. One-third of these miRNAs were intronic with promoters distinct from their host’s, and poor expression correlation was seen between the host gene and the intronic miRNA genes with independent promoters. These novel miRNA promoters also shared other attributes of protein-coding genes: approximately 51% of novel miRNA promoters contain a CpG island, 19% have a TATA box, and 21% have a TFIIB recognition element^262^. Interestingly, it was also discovered that intronic miRNAs with independent transcription initiation regions were located farther from their host gene transcriptional start site^262^ suggesting that these intronic miRNAs evolved to adopt nearby novel independent transcriptional start sites in order to make the transcription of pri-miRNAs more efficient. One example of this possible evolution is the uncoupling of the ancient myosin gene MYH7b and its intronic miRNA miR-499^270^. Other miRNAs whose expression is separate from its host gene expression are miR-199b^271^ and miR-211^272^ which are involved in heart failure and melanoma migration and invasion, respectively, showing that this type of transcriptional regulation of miRNAs can be involved in a spectrum of diseases.

Along with histone modification and chromatin regulation, DNA methylation is also responsible for miRNA transcriptional regulation. miRNA species that show cell type-specific patterns of expression or those that are important in the maintenance of cell identity are prime targets for epigenetic control^273,274^. miR-200c and miR-141 are members of the miR-200 family of miRNAs and have been shown to be important regulators of epithelial-to-mesenchymal transition^275,276^, and dysregulation of miR-200c occurs in multiple types of cancer^147,277^. Recently, it was shown that DNA methylation plays a role in cell type-specific expression of miR-200c and miR-141 and that aberrant DNA methylation of the miR-200c/141 CpG island is closely linked to their silencing in cancer cells^278^. Silencing of miRNAs due to DNA methylation has also been shown to be an early event in pancreatic cancer carcinogenesis^279^, pancreatic cancer development^280^, and pancreatic cancer progression and metastasis^260^.

Aside from epigenetic regulation, traditional transcription factors can also regulate the expression of independently transcribed miRNAs. Since these regulatory proteins preferentially bind nucleosome-free DNA^281,282^—which is found around novel miRNA promoters—nucleosome occupancy can be used to determine transcription factors that regulate miRNA expression. E-box elements recognized by the transcription factor MITF were identified within 1kb upstream and 250 base pairs downstream of 175 novel miRNA transcription initiation sites^262^. Ozsolak et al. (2008) also found c-Myc bound to several MITF-bound miRNA promoters suggesting that transcription factors likely to modulate miRNA expression can be rapidly identified through identification of their DNA binding elements within the nucleosome-depleted regions surrounding novel miRNA promoters. Marson et al. studied the transcriptional regulatory circuit of embryonic stem cells that incorporates both protein-coding and miRNA genes. To identify miRNA promoters, genomic coordinates of H3K4me3-enriched loci from several cell types were used to create a library of transcriptional start sites in both human and mouse^283^. Using their library, they found a major peak of H3K4me3 at the mouse *Slit2* promoter and a minor peak of H3K4me3 downstream of the mouse *Slit2* promoter. This suggested a candidate site for an alternative promoter for mir-218-1 expression.

To further understand the expression, location, and transcriptional regulation of mir-218-1 in pancreatic ductal adenocarcinoma, the level of mir-218-1 was examined in a panel of pancreatic cancer cell lines: HPDE, BxPC-3, Su.86.86, Capan-1, Capan-2, HPAF-II, MIA PaCa-2, PANC-1, and Hs 766T. mir-218-1 expression was variable in pancreatic cancer cell lines and its expression did not correlate with the expression of its host gene *SLIT2*. Chromatin immunoprecipitation was utilized to determine the location and chromatin landscape of the candidate mir-218-1 alternative promoter. Data from Vrba et al. (2011) allowed me to focus on a specific genomic region of *SLIT2* for ChIP analysis. A peak of H3K4me3 enrichment was mapped to a 1kb region within intron 4 of the *SLIT2* gene and enrichment of H4ac overlapped the H3K4me3 peak in all pancreatic cancer cell lines tested. Sequenom analysis for CpG methylation was used to evaluate the methylation status of the candidate mir-218-1 alternative promoter. Although there is no CpG island in this region, isolated CpG sites are present. The candidate mir-218-1 alternative promoter was unmethylated in all cell lines queried upstream of the 1kb region containing the H3K4me3 peak and encompassing the second half of the 1kb region. The third region spanning the first half of the 1kb region was methylated in HPDE and all pancreatic cancer cell lines except PANC-1 and Hs 766T.

TRANScription FACtor database (TRANSFAC) is a database of eukaryotic transcription factors, their DNA binding sites and DNA binding profiles used to predict potential transcription factor binding sites. The database is centered on transcription factors and their DNA binding sites with regard to their structural and functional features extracted from the scientific literature^284^. Binding of a transcription factor to its binding site is documented by determining the location of the site, its sequence and the experimental method applied. All sites that correspond to one transcription factor are aligned and used to construct a position-specific scoring matrix. Inputting the 1kb region into TRANSFAC resulted in a number of predicted transcription factor binding sites including Myc/Max, TCF1/LEF1, TFIIA, Evi-1, and NF-κB. I decided to further examine the potential binding of NF-κB to the mir-218-1 alternative promoter due to its role as a transcriptional activator^285–287^.

Chromatin immunoprecipitation was also utilized to determine the binding of NF-κB to the candidate mir-218-1 alternative promoter. NF-κB was bound to the candidate mir-218-1 alternative promoter in pancreatic cancer cell lines that express mir-218-1 in the absence of *SLIT2* and contain no DNA CpG methylation at the predicted NF-κB binding site. Transfection of pancreatic cancer cell lines that express mir-218-1 in the absence of *SLIT2* with two different oligos to the p65 subunit of NF-κB as well as an inhibitor to IKK, an upstream regulator of NF-κB signaling, led to an increase in pre-mir-218-1 and mir-218-1 expression indicating that NF-κB is a transcriptional repressor of mir-218-1 when expressed from its alternative promoter in pancreatic cancer.

## Materials and methods

### Transfection, treatment, nucleic acid isolation, and qPCR

Pancreatic cancer cell lines Hs 766T and Su.86.86 were plated at a density of 2×10^4^ cells/cm^2^ in individual 6-well tissue culture plates. Twenty-four hours after plating, cells were transfected for 24, 48, or 72 hours with one of two different oligos (siNF-κB 5’-ACAGUAGGAAGAUCUCAUC-3’ and sip65 5’-UUUACGUUUUCUCCUCAAUC-3’) to the p65 subunit of NF-κB at a concentration of 100 nM using Oligofectamine (Life Technologies, Benicia, CA). After transfection, Hs 766T were collected in Trizol (Sigma Aldrich, St. Louis, MO), Su.86.86 were scraped into cell lysates, and processed as described below.

For treatment, Hs 766T were plated at a density of 3×10^4^ cells/cm^2^ in individual 6-well tissue culture plates. Forty-eight hours after plating, cells were treated with 1μM, 2 μM, 5 μM, or 10 μM of BMS-345541 in DMSO (Sigma Aldrich, St. Louis, MO). Twenty-four hours after treatment, cells were collected in Trizol (Sigma Aldrich, St. Louis, MO) and processed as described below.

Both mature and miRNA were extracted together using the Qiagen miRNeasy kit (Qiagen, Valencia, CA) and 1 μg of RNA was reverse transcribed using either 1 μg/ml random primers and Superscript II (Life Technologies, Benicia, CA) or the miSCRIPT II RT kit (Qiagen, Valencia, CA). Primers to *NFKB1* and *18s* were designed using the Roche Universal Probe Library assay design center and qPCR was performed using Quanta PerfeCTa Supermix, Low Rox (Quanta BioScience, Gaithersburg, MD) on an ABI One Step Sequence Detection System (Life Technologies, Benicia, CA). miSCRIPT primer sets to hsa-pre-mir-218-1 (pre-mir-218-1), hsa-mir-218-1 (mir-218-1), hsa-mir-218-2 (mir-218-2), and hsa-U6 snRNA (U6) were purchased from Qiagen (Qiagen, Valencia, CA) and qPCR was performed using the Qiagen miSCRIPT SYBR^®^ Green PCR kit (Qiagen, Valencia, CA) on an ABI One Step Sequence Detection System (Life Technologies, Benicia, CA). Differences in expression between cancer cell lines and HPDE were determined using the comparative Ct method described in the ABI user manual relative to U6 for mir-218-1 and mir-218-2. miRNA was isolated from each cell line in duplicate and each miRNA sample was analyzed by qPCR in triplicate. Differences in expression between siRNA transfected and control non-transfected cells were determined using the comparative Ct method described in the ABI user manual relative to *18s* for *NFKB1* and relative to U6 for pre-mir-218-1 and mir-218-1. Hs 766T was transfected with siRNA in duplicate and each transfection was analyzed by qPCR in triplicate. Differences in expression between BMS-345541 treated and control non-treated cells were determined using the comparative Ct method described in the ABI user manual relative to U6 for pre-mir-218-1 and mir-218-1. Hs 766T was treated in duplicate and each treatment was analyzed by qPCR in triplicate. Primer sequences are listed in Table 3.1.

**Table 3.1:**
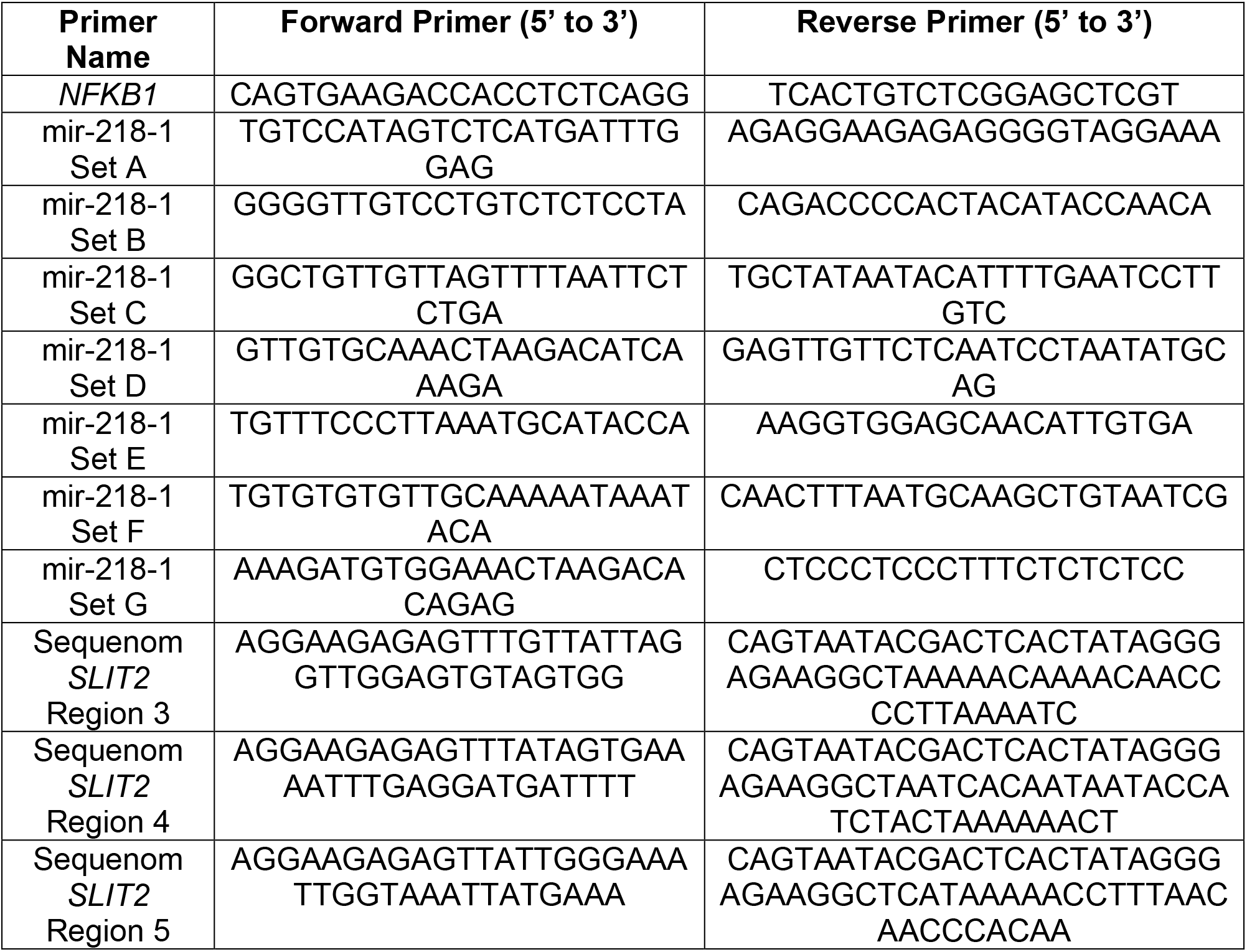
Primer sequences used in Chapter 3. Primer sequences used to determine gene expression, chromatin enrichment of histone modifications, and DNA CpG methylation via Sequenom analysis in Chapter 3. All primers are written from 5, to 3,.

### Chromatin Immunoprecipitation

Cell lysates were collected and processed using the EZ-ChIP kit (Millipore, Billerica, MA). Antibodies against H3K4me3 and NF-κB were purchased from Millipore (#04-745 and MAB3026, respectively). Each cell line was immunoprecipitated twice using the protocol described in Chapter 2. Equal amounts of ChIP and input DNA were used for qPCR analysis. Primers upstream of the 1kb region containing the mir-218-1 alternative promoter (primer sets A and B), downstream of the 1kb region containing the mir-218-1 alternative promoter (primer sets F and G), and encompassing the 1kb region containing the mir-218-1 alternative promoter and NF-κB binding site (primer sets C, D, and E) were designed using the Roche Universal Probe Library assay design center. qPCR was performed using Power SYBR^®^ Green Master Mix (Life Technologies, Benicia, CA) on an ABI Prism 7500 Sequence Detection System (Life Technologies, Benicia, CA). Differences in immunoprecipitation were determined using the fold enrichment method (Life Technologies, Benicia, CA). Each ChIP sample was processed in triplicate. Primer sequences are listed in Table 3.1.

### Sequenom

Sequenom was performed as described in Chapter 2 with region specific primers designed to the alternative mir-218-1 promoter and upstream of the mir-218-1 alternative promoter (Sequenom, San Diego, CA). Primer sequences are listed in Table 3.1.

### Dual-luciferase reporter assay

Dual-Luciferase reporter assays were performed according to the manufacturer’s protocol (Promega, Madison, WI). The 1kb region associated with the mir-218-1 alternative promoter was cloned into pGL3 containing a luciferase reporter gene. HPDE and pancreatic cancer cell lines were plated at a density of 3×10^4^ cells/cm^2^ in individual 6-well tissue culture plates. Forty-eight hours after plating, cells were transiently transfected with pGL3 containing a luciferase reporter gene or pGL3-mir-218-1TSS and a control pTK Renilla reporter construct. Twenty-four hours after transfection, cells were lysed, protein concentration was determined using the BCA method and analyzed on a luminometer. Transcriptional activity was measured for 5 μg of protein as relative light units of luciferase activity to renilla activity. The light units were then normalized to the activity of pTKRenilla control and the results were expressed as fold activation of transcription over HPDE.

### SDS-PAGE and Western Blot

Cell lysates from Su.86.86 were prepared and separated by 7% SDS-PAGE and elecrophoretically transferred to nitrocellulose. Monolayers of cells were washed with calcium and magnesium-free phosphate buffered saline (CMF-PBS) containing 1 mM phenylmethylsulfonyl fluoride (PMSF). Cells were scraped in CMF-PBS, transferred to a microcentrifuge tube, and centrifuged. The pellet was lysed with 2X SDS Sample Buffer (0.25M Tris-HCL pH 6.8, 10% SDS, 25% glycerol) and 30 μg of cellular protein was loaded per lane. Protein concentrations were measured using the BCA method. Antigens were detected by primary antibodies followed with peroxidase-conjugated anti-rabbit IgG. Protein bands were identified by chemiluminescence (NEN, Boston, MA) exposed on X-OMAT AR film (Kodak, Rochester, NY). Images of western blots were captured using Metamorph Version 3.0 (Hollis, NH). The antibodies to NF-κB (SC-372) and beta-tubulin (SC-9104) were purchased from Santa Cruz Biotechnologies (Dallas, TX).

## Results

### mir-218 expression in pancreatic cancer cell lines

Since mir-218-1 and mir-218-2 are encoded within the *SLIT2* and *SLIT3* genes, respectively, expression of these two miRNAs were characterized in a large number of pancreatic cancer cell lines and immortalized human pancreatic ductal epithelial cells. To examine miRNA expression, miRNA was isolated, reverse transcribed, and queried with primers designed to either mir-218-1 or mir-218-2. qPCR analysis indicated that mir-218-1 expression is variable in pancreatic cancer cells and does not correlate with *SLIT2* expression (Figures 3.1A and 2.1B) while mir-218-2 is only expressed in PANC-1 and correlates with *SLIT3* expression (Figures 3.1B and 2.1C). Based on *SLIT2* and mir-218-1 expression, pancreatic cancer cell lines can then be classified into four different subgroups: 1) those that express *SLIT2*, but do not express mir-218-1 (BxPC-3), 2) those that express both *SLIT2* and mir-218-1 (Su.86.86), 3) those that do not express *SLIT2*, but express mir-218-1 (Capan-1, Capan-2, and Hs 766T), and 4) those that do not express *SLIT2* or mir-218-1 (HPAF-II, MIA PaCa-2, and PANC-1). These results suggest that mir-218-1 may be transcribed from a promoter separate from the *SLIT2* promoter in a subset of pancreatic cancer.

**Figure 3.1:**
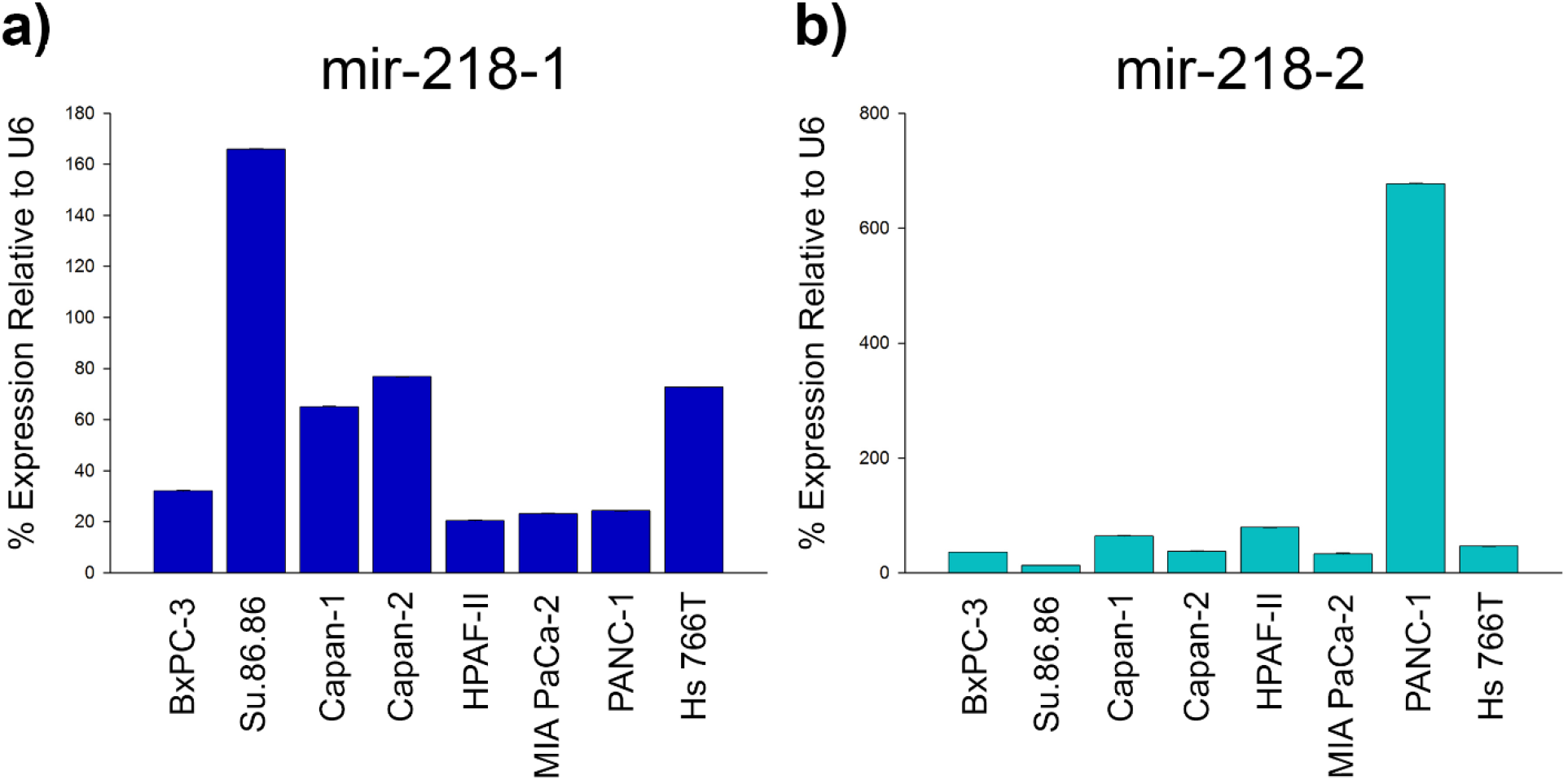
mir-218-1 and mir-218-2 expression in pancreatic ductal adenocarcinoma cell lines. miRNA was extracted with Trizol and cDNA was prepared using the Qiagen miSCRIPT II RT kit. Quantitative PCR (qPCR) was carried out and miRNA levels were normalized to immortalized human pancreatic ductal epithelium relative to U6. Expression of miRNA as detected by qPCR for (a) mir-218-1 and (b) mir-218-2. mir-218-1 expression is variable in pancreatic ductal adenocarcinoma cell lines and does not correlate with *SLIT2* expression while mir-218-2 expression correlates with *SLIT3* expression.

### Characterization and transcriptional activity of a candidate mir-218-1 alternative promoter

It has been shown previously that expression of intronic miRNAs can be uncoupled from the expression of their host genes. Looking at chromatin structures to identify miRNA promoters, Ozsolak et al. found that approximately one-third of intronic miRNAs have transcription initiation regions independent of their host gene promoters^262^. Interestingly, it has also recently been published that a peak of H3K4me3 enrichment exists within the *SLIT2* gene in human mammary fibroblasts and mouse embryonic fibroblasts^283,288^. Since H3K4me3 enrichment is a chromatin mark associated with promoter regions^228,266^, we mapped the peak of H3K4me3 enrichment in human mammary fibroblasts found by Vrba et al. (2013) to a 1kb region within intron 4 of the *SLIT2* gene and performed chromatin immunoprecipitation to determine whether or not this peak exists in pancreatic epithelium. We analyzed immortalized human pancreatic ductal epithelium and several pancreatic cancer cell lines that contain either *SLIT2* mRNA and mir-218-1 expression or only mir-218-1 expression. In all cell lines examined, a peak of H3K4me3 enrichment within intron 4 of the *SLIT2* gene was discovered. The highest chromatin enrichment within the peak was found across primer sets C, D, and E, which correspond to the 1kb region containing H3K4me3 enrichment in human mammary fibroblasts analyzed by Vrba et al. (2013). I also found that H3K4me3 enrichment between the different subtypes of cell lines was also equal to one another (Figure 3.2A). These results suggest that mir-218-1 expression in cell lines that do not express *SLIT2* could result from the activation of a candidate alternative promoter present in both immortalized pancreatic ductal epithelium and pancreatic ductal adenocarcinoma.

**Figure 3.2:**
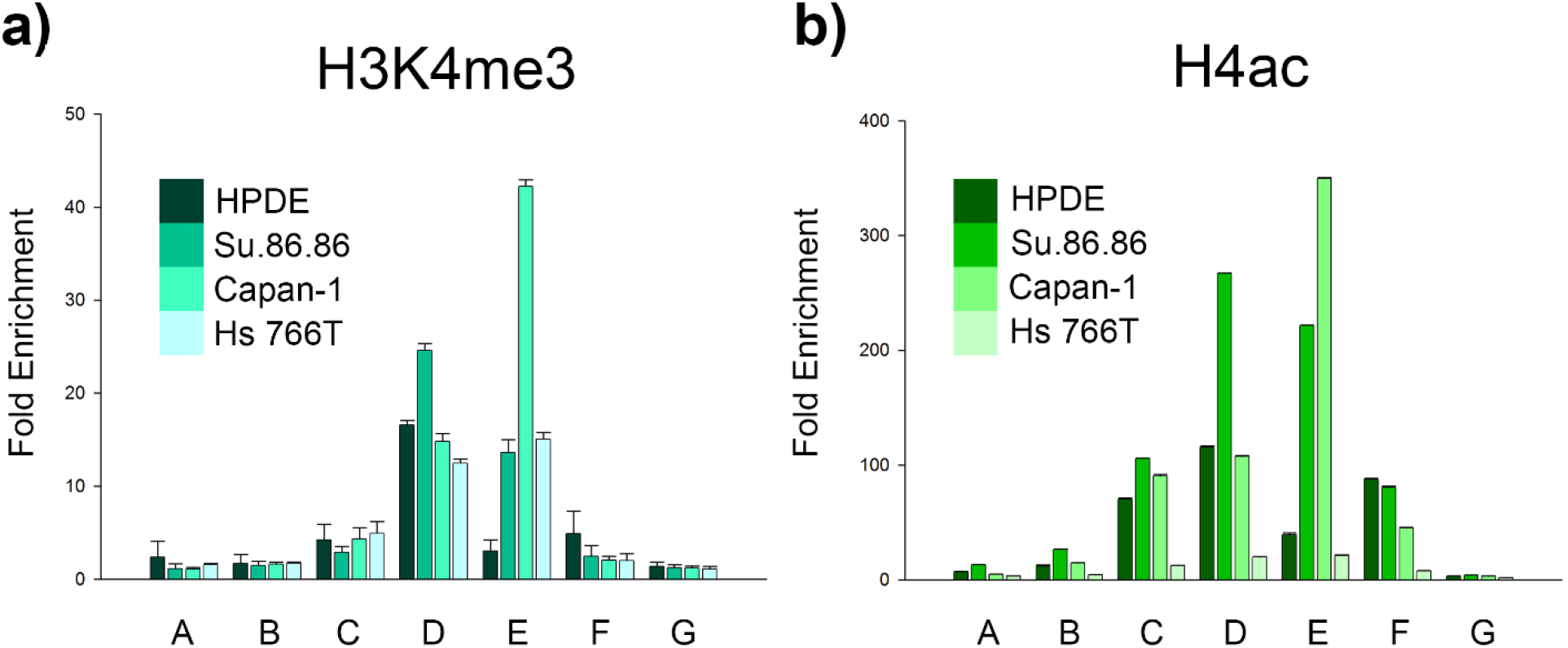
Chromatin enrichment of histone modifications at the putative mir-218-1 alternative promoter. Cell lysates were crosslinked, scraped, sonicated, precleared, and incubated overnight with a rabbit polyclonal antibody to H4ac or a rabbit monoclonal antibody to H3K4me3. 10% of cell lysates were set aside as input DNA prior to preclearing. Immunoprecipitated DNA was washed, crosslinks were reversed, protein was digested, and equal amounts of immunoprecipitated DNA and input DNA were analyzed via quantitative PCR. A difference in immunoprecipitation was determined using the fold enrichment method. Chromatin enrichment at the putative mir-218-1 alternative promoter for (a) H3K4me3 and (b) H4ac. Enrichment for H3K4me3 is approximately equal in all pancreatic cell lines. Enrichment for H4ac overlapped the enrichment for H3K4me3, but did not correlate with mir-218-1 expression.

To determine whether or not transcription from the candidate alternative promoter could be regulated by epigenetic mechanisms, ChIP for H4ac was performed as well as Sequenom analysis for DNA CpG methylation. In all cell lines tested, a peak of H4ac enrichment overlapped the peak of H3K4me3 enrichment with the highest enrichment within the 1kb region itself. Enrichment of H4ac did not correlate with mir-218-1 expression since Capan-1 and Hs 766T express mir-218-1 at the same level, yet enrichment of H4ac in Capan-1 is much higher than that in Hs 766T (Figure 3.2B). While the genomic region analyzed contained CpG sites, they do not form a CpG island. Sequenom analysis determined that the region directly upstream of the 1kb region harboring the mir-218-1 alternative promoter (Region 3) remains relatively unmethylated in HPDE and all pancreatic cancer cell lines queried. The region encompassing the second half of the 1kb region (Region 4) also remains relatively unmethylated with one CpG site showing various amounts of methylation in BxPC-3, Su.86.86, Capan-1, and Capan-2. The region covering the first half of the 1kb region (Region 5); however, is methylated in all cell lines except PANC-1 and Hs 766T (Figure 3.3).

**Figure 3.3:**
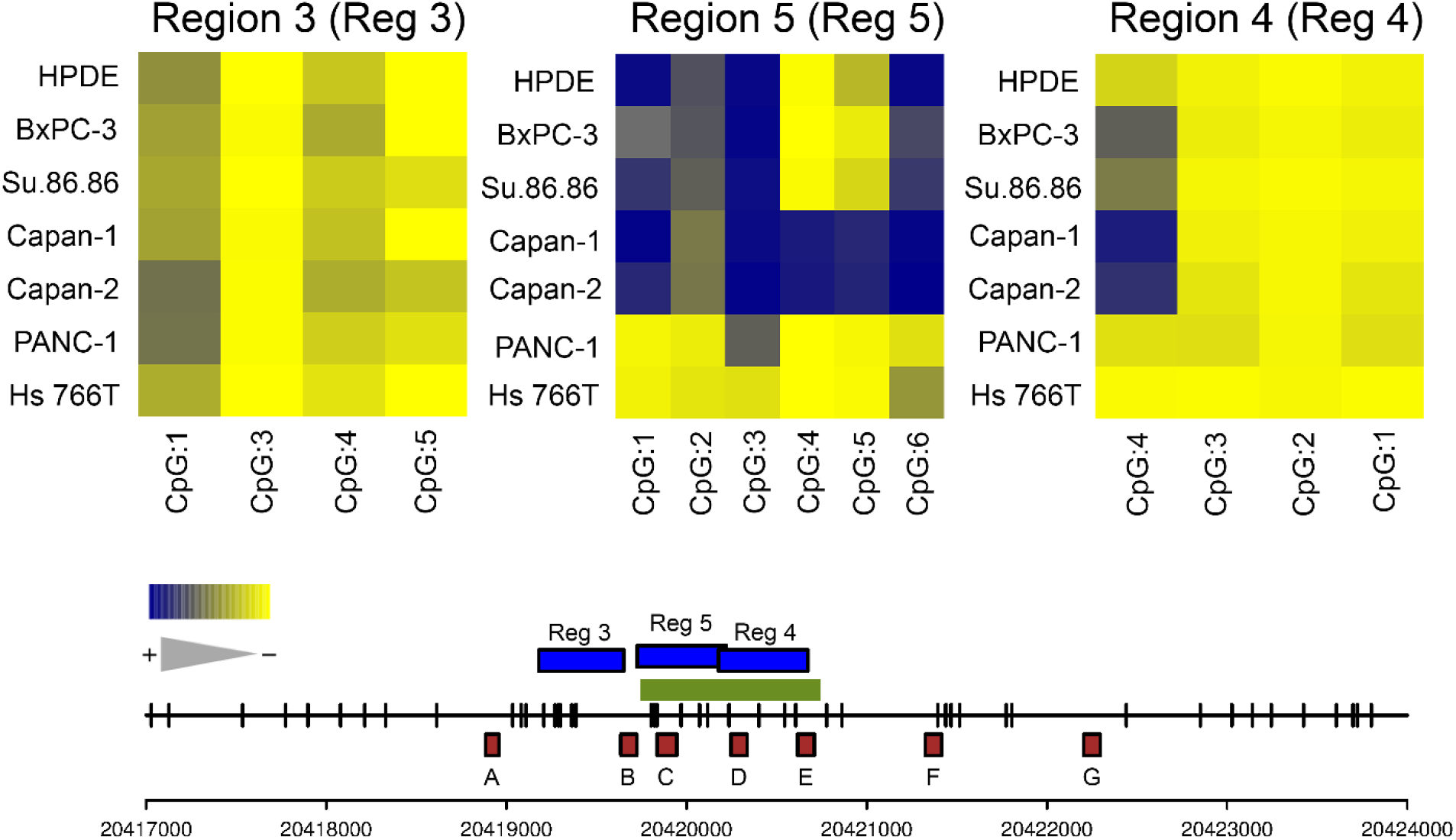
DNA CpG methylation at the putative mir-218-1 alternative promoter. Genomic DNA was isolated from pancreatic ductal adenocarcinoma cell lines and sodium bisulfite converted. Sodium bisulfite converted DNA was then amplified with region-specific PCR primers, *in vitro* transcribed, cleaved with RNaseA, and analyzed using a matrix-assisted laser desorption/ionization-time-of-flight mass spectrometer. Data is presented as the average percent methylation within each fragment. Yellow indicates 0% DNA CpG methylation while blue indicates 100% methylation. The green bar represents the peak of H3K4me3 chromatin enrichment, the blue bars represent the regions of the candidate mir-218-1 alternative promoter analyzed by Sequenom, the red boxes represent the regions analyzed by ChIP, and the numbers along the bottom represent the location in the genome in relation to the UCSC Genome browser hg19 assembly. At the putative mir-218-1 alternative promoter, Region 3 remains relatively unmethylated in HPDE and all pancreatic cancer cell lines queried. Region 4 also remains relatively unmethylated with one CpG site showing various amounts of methylation in BxPC-3, Su.86.86, Capan-1, and Capan-2. Region 5; however, is methylated in all cell lines except PANC-1 and Hs 766T.

Dual-luciferase reporter assays were used to establish transcriptional activity from the 1kb region. BxPC-3 which express *SLIT2* mRNA, but little mir-218-1 shows 3-fold higher transcriptional activity of mir-218-1 than HPDE. Su.86.86, which express both *SLIT2* and mir-218-1, has the highest transcriptional activity of mir-218-1 from the alternative promoter at 4-fold higher than HPDE. PANC-1, which does not express *SLIT2* or mir-218-1, displays 3-fold enrichment of mir-218-1 transcriptional activity over HPDE. Finally, Hs 766T which does not express *SLIT2*, but does express mir-218-1 also shows a 3-fold increase in transcriptional activation over HPDE (Figure 3.4). These results indicate that the chromatin in this area of the genome is open, contains epigenetic marks permissible for transcription, and contains transcriptional activity even in cell lines that do not express high levels of mir-218-1.

**Figure 3.4:**
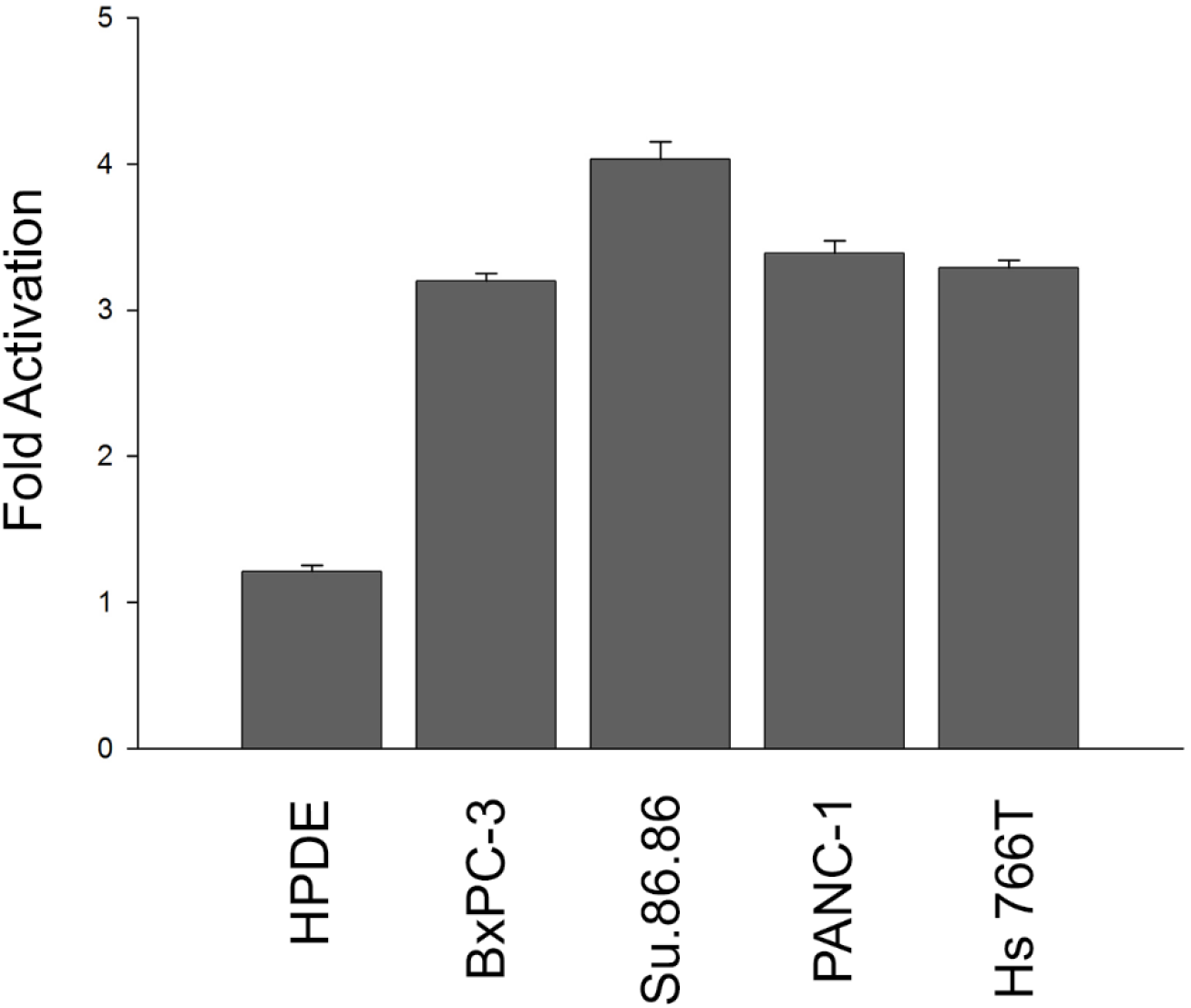
Transcriptional activity at the putative mir-218-1 alternative promoter. The 1kb region containing the putative mir-218-1 alternative promoter was cloned into the pGL3 vector containing a luciferase reporter gene. Twenty-four hours after transfection the cells were lysed, protein concentration was determined, and 5 μg of protein was analyzed on a luminometer. Transcriptional activity was measured as relative light units of luciferase activity to renilla activity. The relative light units were then normalized to the activity of pTKRenilla control and expressed as fold activation of transcription over human pancreatic ductal epithelium. The putative mir-218-1 alternative promoter contains transcriptional activity even in pancreatic ductal adenocarcinoma cells that do not contain mir-218-1 expression suggesting the presence of alternative regulatory elements outside of this 1kb region.

### Transcriptional regulation of the candidate mir-218-1 alternative promoter by NF-κB

Since the 1kb region containing the candidate mir-218-1 alternative promoter shows transcriptional activity in all pancreatic cancer cell lines, we decided to probe this region for transcription factor DNA binding sites. To identify these sites, the 1kb region was input into a computer program (the TRANScription FACtor database—TRANSFAC)^284^ to predict potential transcription factor binding sites. The program predicted a NF-κB DNA binding site within the 1kb region. We decided to further examine the potential role of NF-κB binding at the candidate mir-218-1 alternative promoter due to its known activity as a transcriptional activator^285–287^ and role in pancreatic cancer progression^289,290^. Upon doing so, we came upon a putative NF-κB binding site located in the middle of ChIP primer set D, which is located in the middle of the 1kb region. Therefore, we performed ChIP to determine if NF-κB binds to the candidate mir-218-1 alternative promoter. To do this, we used HPDE and Su.86.86, which express both *SLIT2* and mir-218-1 as well as Capan-2 and Hs 766T which only express mir-218-1. HPDE showed no enrichment of NF-κB over the no antibody control at primer sets C and D, but showed 3-fold enrichment at primer set E. Su.86.86 contains no enrichment of NF-κB at any of the primer sets. Capan-2 showed very little enrichment of NF-κB at primer sets C and D, but showed 6-fold enrichment at primer set E. Hs 766T showed little-to-no enrichment of NF-κB at primer sets C and E, but contained 10-fold enrichment at primer set D containing the putative NF-κB binding site (Figure 3.5). These results indicate that NF-κB binds to a predicted cis regulatory element in the mir-218-1 candidate alternative promoter in cell lines that do not have DNA methylation at the NF-κB binding site.

**Figure 3.5:**
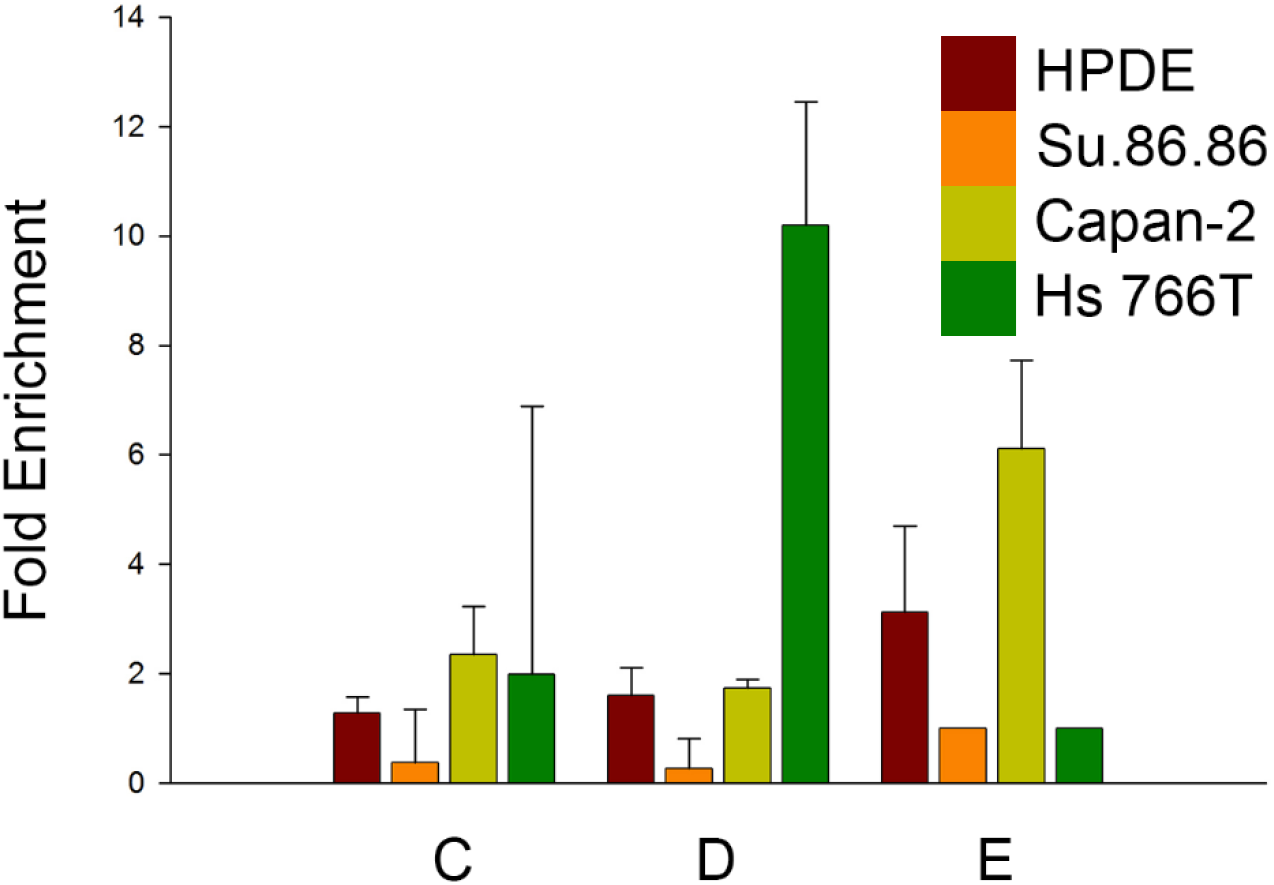
Chromatin enrichment of NF-κB at the putative mir-218-1 alternative promoter. Cell lysates were crosslinked, scraped, sonicated, precleared, and incubated overnight with a mouse monoclonal antibody to NF-κB. 10% of cell lysates were set aside as input DNA prior to preclearing. Immunoprecipitated DNA was washed, crosslinks were reversed, protein was digested, and equal amounts of immunoprecipitated DNA and input DNA were analyzed via quantitative PCR. A difference in immunoprecipitation was determined using the fold enrichment method. NF-κB binds to a predicted cis regulatory element in the mir-218-1 candidate alternative promoter in cell lines that do not have DNA methylation at the predicted NF-κB binding site.

To further determine the role of NF-κB on mir-218-1, we wanted to determine whether or not NF-κB controls mir-218-1 expression. For this purpose, we used previously designed siRNA oligos to the p65 subunit of NF-κB (siNF-κB and sip65) that have been previously validated. Transfection of Hs 766T with siNF-κB led to a time-dependent 80% decrease in *NFKB1* expression that was greatest at 24 hours and an increase in pre-mir-218-1 at 72 hours and an increase in mir-218-1 at 24 hours (Figure 3.6A). Transfection of Hs 766T with sip65 led to a time-dependent 90% decrease in *NFKB1* that was greatest at 48 hours and an increase in both pre-mir-218-1 and mir-218-1 at 48 hours (Figure 3.6B). Transfection of Su.86.86 with siNF-kB also led to a time-dependent decrease in NF-κB protein (Figure 3.6C). These results indicate that NF-κB is acting as a transcriptional repressor of mir-218-1 when expressed from its alternative promoter in pancreatic cancer. While NF-κB is generally known as a transcriptional activator, several studies indicate that it can also function as a repressor^291–293^.

**Figure 3.6:**
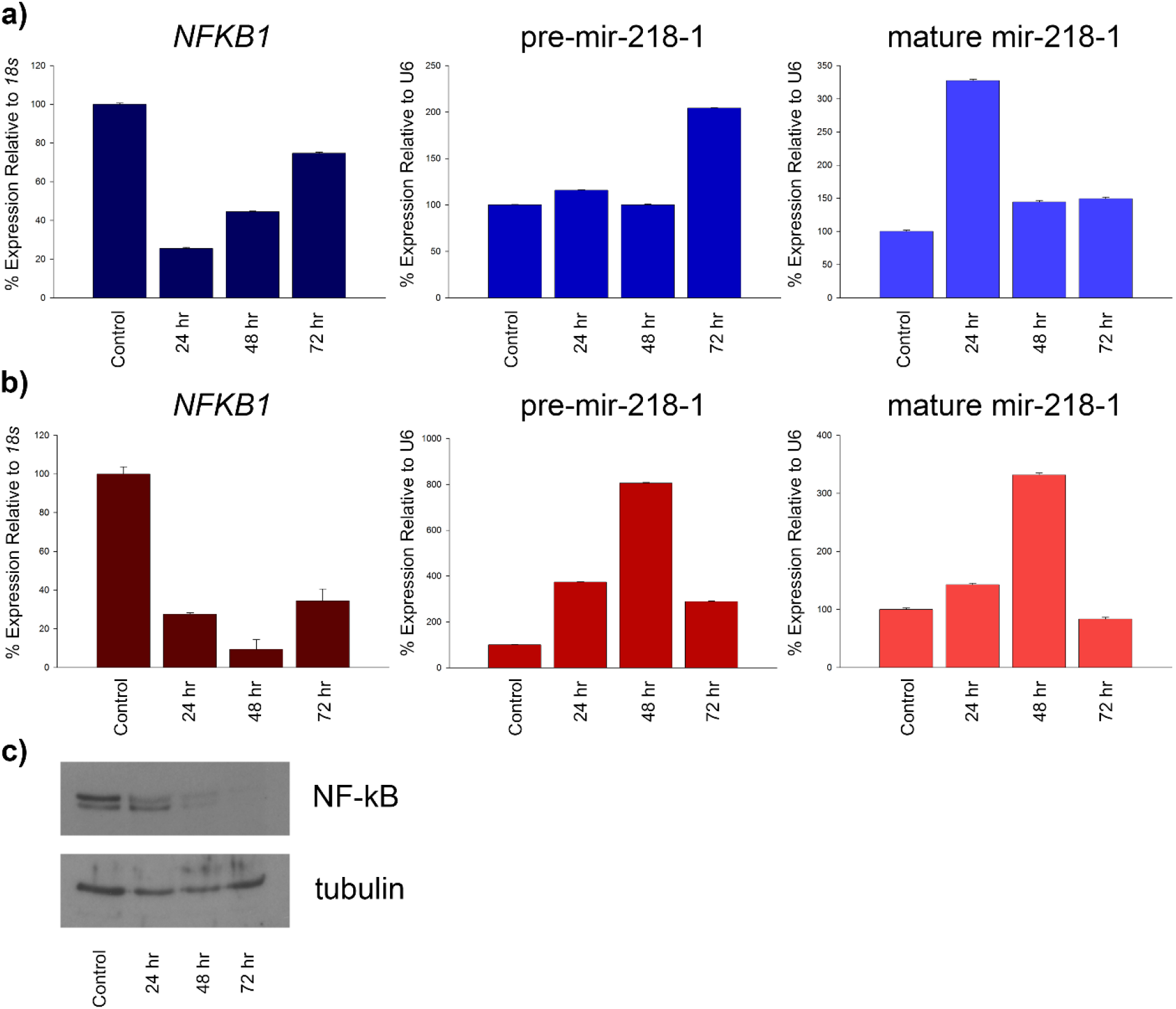
NF-κB is a transcriptional repressor of both pre-mir-218-1 and mature mir-218-1. Hs 766T were transfected with 100 nM of siNF-κB or sip65 for 24, 48, or 72 hours. After transfection, cells were collected in Trizol, mRNA and miRNA was isolated, and reverse transcribed. Quantitative PCR (qPCR) was performed using intron-spanning primers designed by the Roche Universal Probe Library Assay Design Center or using primers designed by the Qiagen miScript system. Differences in expression between siRNA-transfected and control non-transfected cells were determined using the comparative Ct method relative to *18s* or U6. Expression of *NFKB1*, pre-mir-218-1, or mir-218-1 as detected by qPCR for (a) siNF-κB or (b) sip65. Su.86.86 were transfected with 100 nM of siNF-κB for 24, 48, or 72 hours. After transfection, total cell lysates were extracted in 2X SDS-sample buffer, protein concentration was determined, and 30 μg of protein per lane was analyzed by SDS-PAGE gel. After electrophoresis, proteins were transferred to nitrocellulose membranes and the filters were treated with rabbit polyclonal antibodies to the p65 subunit of NF-κB or beta-tubulin. Immunoblots were developed with a chemiluminescence detection reagent. (c) Expression of NF-κB after transfection with siRNA to the p65 subunit of NF-κB. NF-κB acts as a transcriptional repressor of mir-218-1 when it is expressed from its putative alternative promoter.

To further support the role for NF-κB in transcriptional repression of mir-218-1, we treated Hs 766T with an inhibitor of IκB kinase (IKK), an upstream effector of the NF-κB signaling cascade^294^. The IκBα protein is responsible for keeping NF-κB in its inactive state by binding to NF-κB and masking its nuclear localization sequence^295^. IKK is then responsible for phosphorylating IκBα resulting in the dissociation between IκBα and NF-κB allowing NF-κB to migrate into the nucleus to regulate transcription^296^. Treatment of Hs 766T with IKK inhibitor BMS-345541 led to a dose-dependent increase in pre-mir-218-1 expression ending with a 9-fold increase with the addition of 10 μM of BMS-345541 for 24 hours. There was no increase in mir-218-1 expression in Hs 766T after treatment with BMS-345541 (Figure 3.7). These results further indicate that NF-κB is a transcriptional repressor of mir-218-1 and does not have an effect on miRNA processing.

**Figure 3.7:**
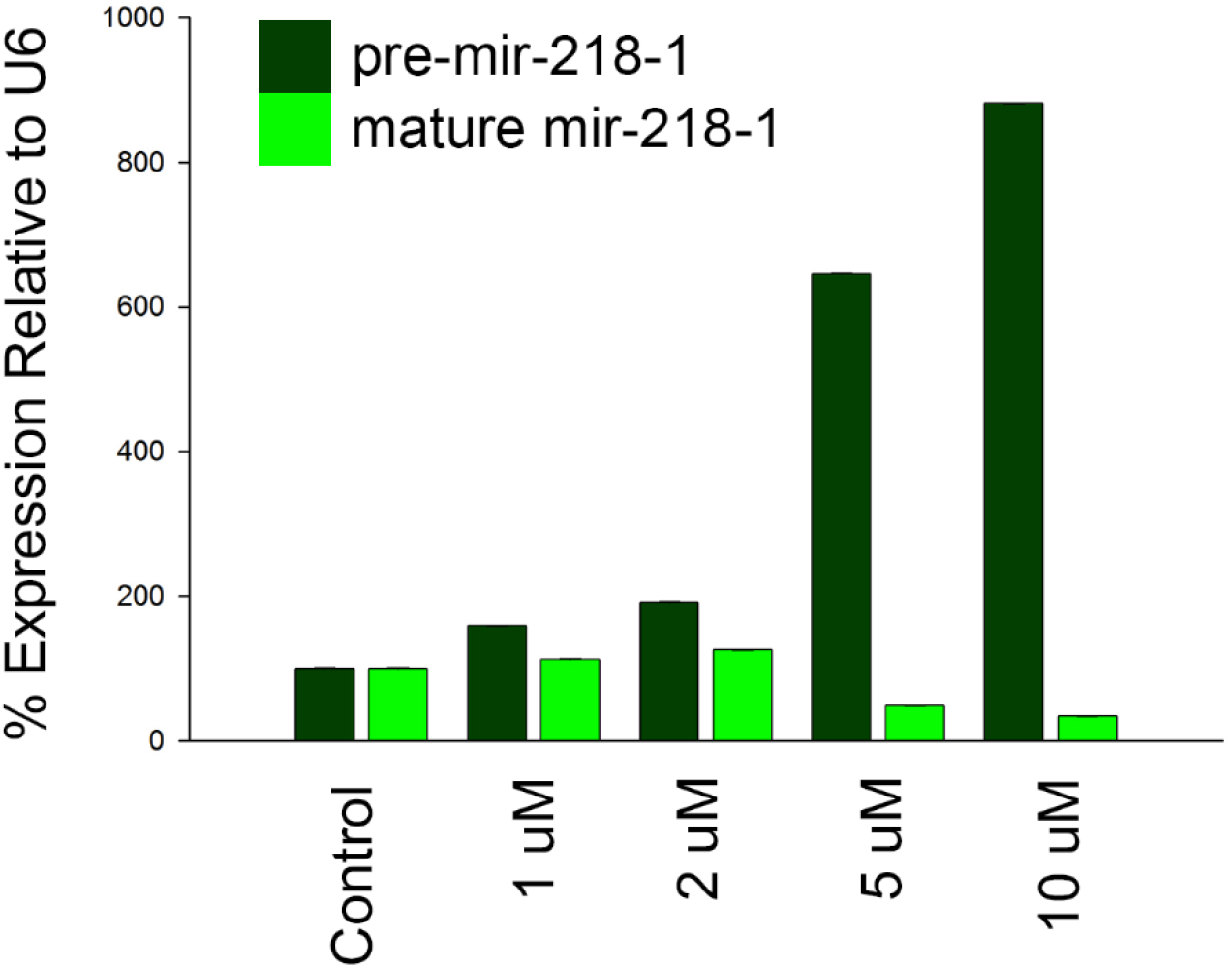
NF-κB signaling is a transcriptional repressor of pre-mir-218-1. Cells were treated with 1 μM, 2 μM, 5 μM, or 10 μM of IKK inhibitor BMS-345541. Twenty-four hours after treatment, cells were collected in Trizol, miRNA was isolated, and reverse transcribed. Quantitative PCR (qPCR) was performed using primers designed by the Qiagen miScript system. Differences in expression between BMS-345541 treated and control non-treated cells were determined using the comparative Ct method relative to U6. Expression of pre-mir-218-1 or mir-218-1 as detected by qPCR. NF-κB signaling acts as a transcriptional repressor of pre-mir-218-1.

## Discussion

Mature miRNAs are 22-24 nucleotide long non-coding RNAs that are evolutionarily conserved and involved in processes including development, tissue homeostasis and cancer. Approximately 50-80% of miRNAs reside within introns of host mRNA genes and thought to be transcriptionally linked with expression of the host gene^124,297^. mir-218-1 and mir-218-2 are intronic miRNAs encoded within the *SLIT2* and *SLIT3* genes, respectively^179^. In this study, we measured the expression of both mir-218-1 and mir-218-2 in pancreatic cancer cell lines and found that expression of mir-218-2 correlates with *SLIT3* expression while expression of mir-218-1 does not correlate with *SLIT2* expression. Therefore, we hypothesized that mir-218-1 has an alternative promoter and transcriptional start site separate from *SLIT2*. Recently it was published that a small peak of chromatin enrichment for H3K4me3 can be found within the *Slit2* gene downstream of the *Slit2* promoter in mouse embryonic fibroblasts^283^. Since H3K4me3 is a validated histone modification associated with active promoter regions^266^, we used the data from Vrba et al. (2013) and mapped the region of H3K4me3 chromatin enrichment in HMFs to a 1kb region in intron 4 of the *SLIT2* gene and performed chromatin immunoprecipitation to survey the histone marks in this area of the genome in pancreatic cancer cell lines. We found that HPDE and all pancreatic cancer cell lines tested contained a peak of both H3K4me3 and H4ac enrichment within the 1kb region of intron 4 of the *SLIT2* gene suggesting that mir-218-1 could have its own novel promoter and transcriptional start site separate from *SLIT2* that may be active in pancreatic ductal adenocarcinoma.

This is not the first example of an intronic miRNA whose expression is uncoupled from its host gene expression. Ozsolak et al. (2008) performed a study looking at chromatin structures to identify miRNA promoters and found that approximately one-third of intronic miRNAs have transcription initiation units independent of their host gene promoters. This group showed that the novel miR-17 transcriptional start site lies approximately 2kb downstream from the host gene transcriptional start site, and, using luciferase assays, elucidated that both the host gene and novel transcription initiation region have promoter activities. Thus, in this case, transcripts emanating from both the host gene transcriptional start site and its novel transcriptional start site may encode miR-17. In our studies, the amount of enrichment of H3K4me3 was approximately equal in all pancreatic cancer cell lines regardless of *SLIT2* and mir-218-1 expression. Additionally, transcriptional activity was recorded from the 1kb region in all pancreatic cancer cell lines using luciferase assays and not in immortalized human pancreatic ductal epithelium. This suggests that the candidate alternative promoter for mir-218-1 may be the only region from which mir-218-1 is transcribed and perhaps the differences in mir-218-1 expression in pancreatic ductal adenocarcinoma depends on other nuclear environmental factors such as epigenetic regulation and the binding of specific transcription factors to the novel mir-218-1 promoter.

While cytosine methylation has been predominately associated with gene silencing, it has also been shown that location of cytosine methylation within the genomic sequence can be key in DNA methylation-induced gene silencing^56–58^. In highly expressed genes, a pattern of low DNA methylation was seen around the promoter; however, the gene body saw a considerable amount of DNA methylation^298^. Sequenom analysis of the 1kb region of the mir-218-1 alternative promoter determined that the region directly upstream of the 1kb region remained unmethylated in all pancreatic cancer cell lines while the first half of the 1kb region was methylated at isolated CpG sites in HPDE and all pancreatic cancer cell lines except PANC-1 and Hs 766T. This suggests that although DNA methylation is present at the candidate mir-218-1 alternative promoter and transcriptional start site, it is unlikely that DNA methylation plays a major role in mir-218-1 expression due to the fact that a CpG island does not exist in this region of the genome. Therefore, the methylation must play an alternative role in transcriptional regulation of mir-218-1, possibly through the regulation of transcription factor binding.

Regulation of gene expression is a central element to cell physiology and is the key for cells to adapt to environmental, mechanical, and chemical stressors. The transcription factor NF-κB plays an important role in regulating the expression of many different genes within the cell. The NF-κB family of transcription factors consists of five members: p50, p52, p65 (RelA), c-Rel, and RelB^294^. It has been shown previously that the RelA member of the NF-κB family is constitutively activated in approximately 67% of primary pancreatic ductal adenocarcinomas and roughly 82% of pancreatic cancer cell lines^299^. Therefore, we transiently transfected KRAS-dependent Su.86.86 and KRAS-independent Hs 766T with two different oligos to the p65 (RelA) subunit of NF-κB. Transfection with siNF-κB in Hs 766T led to a 80% decrease in *NFKB1* mRNA while transfection with sip65 led to a 90% decrease in *NFKB1* expression. Both oligos also gave a coordinate increase in expression of both pre-mir-218-1 and mir-218-1. Transfection with siNF-κB in Su.86.86 led to a time-dependent decrease in NF-κB protein expression. Treatment of Hs 766T with an inhibitor to IκB kinase, an upstream effector of the NF-κB signaling cascade, also led to an increase in pre-mir-218-1 expression suggesting that NF-κB acts as a transcriptional repressor of mir-218-1 when transcribed from its alternative promoter which has been shown previously^291^.

NF-κB dimers bind to κB sites within the promoters of genes and regulate transcription via the recruitment of coactivators and corepressors. Most κB sites show little to no selectivity for a specific NF-κB dimer^300,301^. Furthermore, different NF-κB dimers can be exchanged at the same gene promoter during the same NF-κB response^302^. Since each dimer supports a different transcriptional response^302–304^, this exchange of subunits could be exploited to finely tune the NF-κB transcriptional response or shut it down leading to transcriptional inactivation^302^. Interestingly, nucleosome positioning only marginally reduces the ability of NF-κB to bind to its κB sites^305^. Several different transcription factor prediction algorithms postulated that a κB site existed in the middle of the 1kb region containing the candidate mir-218-1 alternative promoter. Chromatin immunoprecipitation using an antibody to the p65 subunit of NF-κB showed that NF-κB binds to the candidate mir-218-1 alternative promoter in cell lines that do not have DNA methylation at the κB site. This suggests that the DNA methylation found around the κB site may block NF-κB from binding to and repressing mir-218-1 expression from its alternative promoter.

My work suggests that mir-218-1 is transcribed from a novel and independent transcriptional start site located within intron 4 of the *SLIT2* gene in pancreatic ductal adenocarcinoma. It is likely that the enrichment of H3K4me3 and H4ac lead to the transcription of this miRNA from its novel transcription initiation region (Figure 3.8). Inhibition of NF-κB either through transient transfection of siRNA to the p65 subunit or inhibition of an upstream signaling molecule led to an increase in both pre-mir-218-1 and mir-218-1 expression. The p65 subunit also bound directly to the putative κB site within the mir-218-1 alternative promoter in pancreatic cancer cell lines that do not have any DNA methylation in that region. Future directions would be to determine the corepressors that are recruited to the mir-218-1 alternative promoter by NF-κB to promote transcriptional repression of mir-218-1. Additionally, elucidating the correlation between Kras activation, NF-κB activation, and mir-218-1 expression and their potential effect on pancreatic ductal adenocarcinoma invasion would greatly improve the knowledge of the complexity of this disease.

**Figure 3.8:**
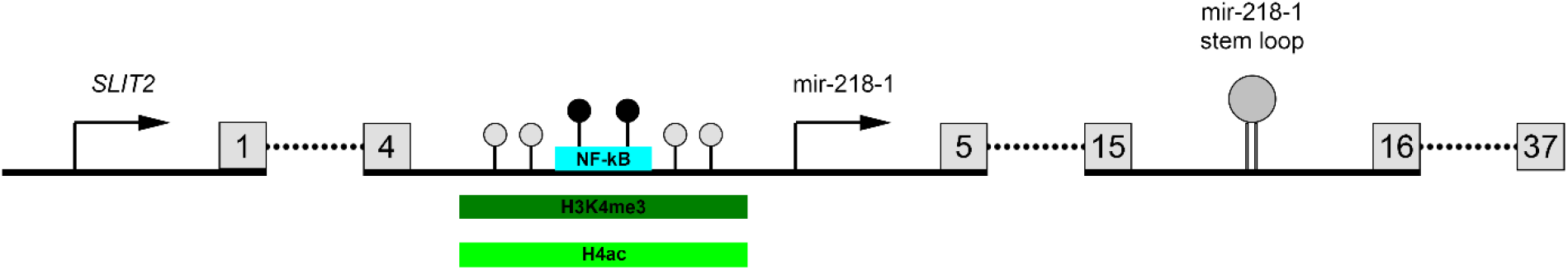
Transcriptional control of the putative mir-218-1 alternative promoter. Transcription of mir-218-1 occurs from a novel and independent promoter and transcriptional start site located within intron 4 of the *SLIT2* gene. Enrichment of H3K4me3 and H4ac leads to transcriptional activation while binding of NF-κB to the alternative promoter in pancreatic ductal adenocarcinoma cell lines with no gene body methylation leads to transcriptional repression. Light gray circles indicate unmethylated CpG sites while black circles indicate methylated CpG sites.

